# Viral Landscape of Gastric Adenocarcinoma Reveals Clinically Relevant Viruses

**DOI:** 10.1101/2025.09.05.673529

**Authors:** Diego Pereira, Fabiano Cordeiro Moreira, Daniel de Souza Avelar, Sérgio Augusto Antunes Ramos, Valéria Cristiane Santos da Silva, Jéssica Manoelli Costa da Silva, Ronald Matheus da Silva Mourão, Rubem Ferreira da Silva, Kauê Sant’Ana Pereira Guimarães, Juliana Barreto Albuquerque Pinto, Marcos da Conceição, Williams Fernandes Barra, Samia Demachki, Samir Mansour Casseb, Rommel Mario Rodriguez Burbano, Paulo Pimentel de Assumpção

**Affiliations:** Graduate Program in Genetics and Molecular Biology, Federal University of Pará, Belém, Pará, Brazil; Oncology Research Center, Federal University of Pará, Belém, Pará, Brazil; Graduate Program in Biotechnology, Federal University of Pará, Belém, Pará, Brazil; 4Graduate Program in Oncology and Medical Sciences, Federal University of Pará, Belém, Pará, Brazil; Graduation in Biotechnology, Federal University of Pará, Belém, Pará, Brazil; Molecular Biology Laboratory, Ophir Loyola Hospital, Belém, Pará, Brazil

**Keywords:** Gastric cancer, virome, human viruses, bacteriophages, viral diversity

## Abstract

**Background:** Gastric cancer (GC) ranks among the most common and lethal cancers worldwide, with poor prognosis mainly due to late diagnosis. Accumulating evidence highlights the role of the gastric microbiome in carcinogenesis through inflammation, genomic instability, and immune modulation. Unlike the bacterial component, the gastric virome remains largely unexplored despite its potential contribution to tumor development. The present study aimed to characterize the virome present in tumor and peritumoral tissues from patients with GC.

**Materials and Methods:** 105 tumor and 85 peritumoral gastric tissues were analyzed. RNA was extracted, libraries were prepared, and sequencing was performed on the Illumina NextSeq 500. Viral reads were classified with Kraken2. Taxonomic profiles, viral abundance and diversity metrics were computed in R, with group differences assessed by Wilcoxon tests and PERMANOVA (p < 0.05).

**Results:** In this study, 38 viral orders and 329 viral genera were identified in gastric tumor and peritumoral tissues. Tumor tissues harbored 210 viral genera, including 109 exclusive to this microenvironment, with bacteriophages comprising the majority, alongside human-infecting and other eukaryotic viruses. *Lymphocryptovirus*, *Cytomegalovirus,* and *Alphapapillomavirus* were enriched in tumors. Alpha and beta diversity analyses revealed no significant differences between tumor and peritumoral tissues, indicating comparable viral richness and composition.

**Conclusion:** These findings underscore the complexity of the gastric virome and provide a foundation for future investigations into the interactions and mechanisms through which the viral community could influence the development of gastric cancer, highlighting its potential role in gastric health and disease.

## 1. Introduction

Gastric cancer (GC) is a malignant neoplasm of the gastrointestinal tract and ranks as the fifth most common cancer worldwide. Although incidence and mortality rates have declined in recent years, GC still exhibits a high mortality rate, reaching approximately 75% in many countries^1, 2^. The low rate of early diagnosis results in most patients being identified at advanced stages of the disease, thereby missing the optimal window for clinical intervention. Under these circumstances, the prognosis remains poor, with a five-year post-surgical survival rate below 30%^3, 4^.

GC is a highly heterogeneous neoplasm, characterized by multiple distinct phenotypes, with most cases being gastric adenocarcinomas (GA). Although often considered a single entity, GC can be classified into two categories based on anatomical location: cardia gastric cancer (upper stomach) and non-cardia gastric cancer (lower stomach)^5^. The main risk factors for cardia GC include obesity and gastroesophageal reflux disease, whereas chronic *Helicobacter pylori* infection is considered the primary cause of non-cardia GC^6, 7^.

In recent decades, several studies have investigated the mechanisms of gastric carcinogenesis using sequencing technologies, with a focus on the relationship between the gastric microbiome and GC development. Recent evidence suggests that microorganisms may play a significant role both in tumorigenesis and in cancer therapeutic strategies^8^. Moreover, the application of high-throughput sequencing technologies in microbiology has revealed that the stomach, previously considered an inhospitable environment for microorganisms, harbors a diverse and functional microbiota^9^.

The mechanisms by which the gastric microbiome contributes to carcinogenesis are not yet fully understood. However, it is recognized that a dysbiotic microbiota can increase the risk of GC through the production of microbial metabolites capable of inducing chronic inflammation in gastric tissue. This inflammatory process promotes the accumulation of mutations and genomic instability in the host, accelerating DNA replication rates and impairing DNA damage repair mechanisms^10, 11^. Moreover, these alterations may be further amplified by interactions between the gastric microbiota, *H. pylori*, and the host immune responses^12–14^.

The human virome is predominantly composed of eukaryotic viruses and bacteriophages, whose dysbiosis has been associated with a variety of diseases, including cancer^15^. Among eukaryotic viruses, oncoviruses are particularly notable for their well-established oncogenic mechanisms. Conversely, bacteriophages can indirectly influence carcinogenesis by modulating the composition and virulence of the bacterial microbiota^16^. Despite advances in characterizing the bacterial component of the gastric tumor microbiome and its association with carcinogenesis, the viral component remains poorly explored, mainly due to technical challenges in detecting viral material and limitations of existing databases^17^.

Therefore, characterization of the virome in GC may provide insights into this neglected component and reveal its potential as a biomarker for diagnosis, prognosis, and novel therapeutic strategies. Accordingly, the present study aimed to characterize the virome present in tumor and peritumoral tissues from patients with GC.

## 2. Materials and Methods

### 2.1. Ethics statement

The study was approved by the Ethics and Research Committee of João de Barros Barreto University Hospital (approval number: 47580121.9.0000.5634) and conducted in accordance with the Declaration of Helsinki. Recruitment and sample collection took place from July 2, 2022, to July 6, 2023. All participants provided written informed consent prior to inclusion.

### 2.2. Sample collection and total RNA extraction

A total of 188 samples, including 103 tumor and 85 peritumoral tissues were collected from patients with gastric adenocarcinoma undergoing surgical resection. From each specimen, approximately 30 mg of tumor tissue was macerated, and RNA was extracted using the TRIzol Reagent method (Thermo Fisher Scientific) in accordance with the manufacturer’s protocol. The integrity and concentration (ng/μL) of the extracted RNA were evaluated using a Qubit 4 Fluorometer (Thermo Fisher Scientific). Only samples with an RNA Integrity Number (RIN) ≥ 5 were considered acceptable.

### 2.3. Library construction and RNA sequencing

RNA libraries were prepared using the TruSeq Stranded Total RNA Library Prep Kit with Ribo-Zero Gold (Illumina, US) following the manufacturer’s protocol, with approximately 1 μg of total RNA per sample in a final volume of 11 μL. Library quality was evaluated using the 2200 TapeStation System (Agilent Technologies, Basel, Switzerland). Sequencing was carried out in paired-end mode on the NextSeq 500 platform (Illumina, US) with the NextSeq 500 MID Output V2 kit (150 cycles), according to the manufacturer’s instructions.

### 2.4. Sequence data pre-processing

Post-sequencing, raw RNA-Seq reads underwent quality control. Read quality was first assessed with FastQC, followed by adapter removal and filtering of low-quality reads using Trimmomatic (v.0.39), applying a quality score threshold of quality value (QV) > 15.

### 2.5. Taxonomic classification

For viral identification, high quality and clean reads were processed with Kraken2 (v2.1.6)^18^, employing the standard database supplemented with the Reference Viral Database (RVDB) (v28.0)^19^ to improve viral detection. The Kraken2 outputs were processed in RStudio software (v2024.04.2) using the Pavian package (v1.2.0).

### 2.6. Viral abundance and composition

The output files were further processed in RStudio using the phyloseq (v1.53.0) and microbiome (v1.19.0) packages to quantify relative abundances at order and genus taxonomic levels. Shared and unique viral genera between groups were assessed and visualized using a Venn diagram with the ggvenn package (v0.1.10). To analyze the virome composition, the identified viral genera were classified according to their predominant host type, enabling a comprehensive evaluation of their distribution across the samples. Differential abundance analysis of viral genera between tumor and peritumoral tissues was performed using the Wilcoxon rank-sum test, with statistical significance set at p < 0.05.

### 2.7. Diversity analysis

Alpha diversity across tissues was assessed using the phyloseq package (v1.53.0), including Observed richness, Shannon, and Simpson indexes. Beta diversity was evaluated with the vegan package (v2.7.1) using Bray–Curtis dissimilarity to measure variation in viral composition among samples. Principal Coordinates Analysis (PCoA) based on Bray–Curtis dissimilarity was also performed to visualize compositional patterns. All boxplots were generated using the ggplot2 package (v3.3.6).

### 2.8. Statistical analysis

Statistical analyses were conducted using PERMANOVA to quantify multivariate community-level differences in viral composition among groups, and the Mann– Whitney (Wilcoxon) test for comparison between two groups. Statistical significance was defined as p-value < 0.05.

## 3. Results

### 3.1. Taxonomic profile of the virome

The microbial taxonomic profile of the 188 gastric tissue samples was characterized using the Kraken2 algorithm. Microbial reads were detected in all samples and classified into bacterial, viral, fungal, protozoan, and archaeal taxa, with only viral taxa retained for downstream analyses. The viral composition was then further examined across different taxonomic levels.

At the order level, 38 viral orders were identified, with *Herpesvirales*, *Lefavirales*, *Bunyavirales*, *Crassvirales*, and *Ortervirales* being the most abundant across the samples overall (Figure 1A). At the genus level, a total of 329 unique viral genera were identified in the gastric tissues. In tumor tissue, *Jouyvirus, Lymphocryptovirus, Betabaculovirus, Cytomegalovirus*, and *Pahexavirus* exhibited the highest abundances (Figure 1B), while in peritumoral tissue, the most abundant genera were *Jouyvirus, Betabaculovirus, Pahexavirus, Lymphocryptovirus*, and Orthotospovirus (Figure 1C).

**Figure 1.**
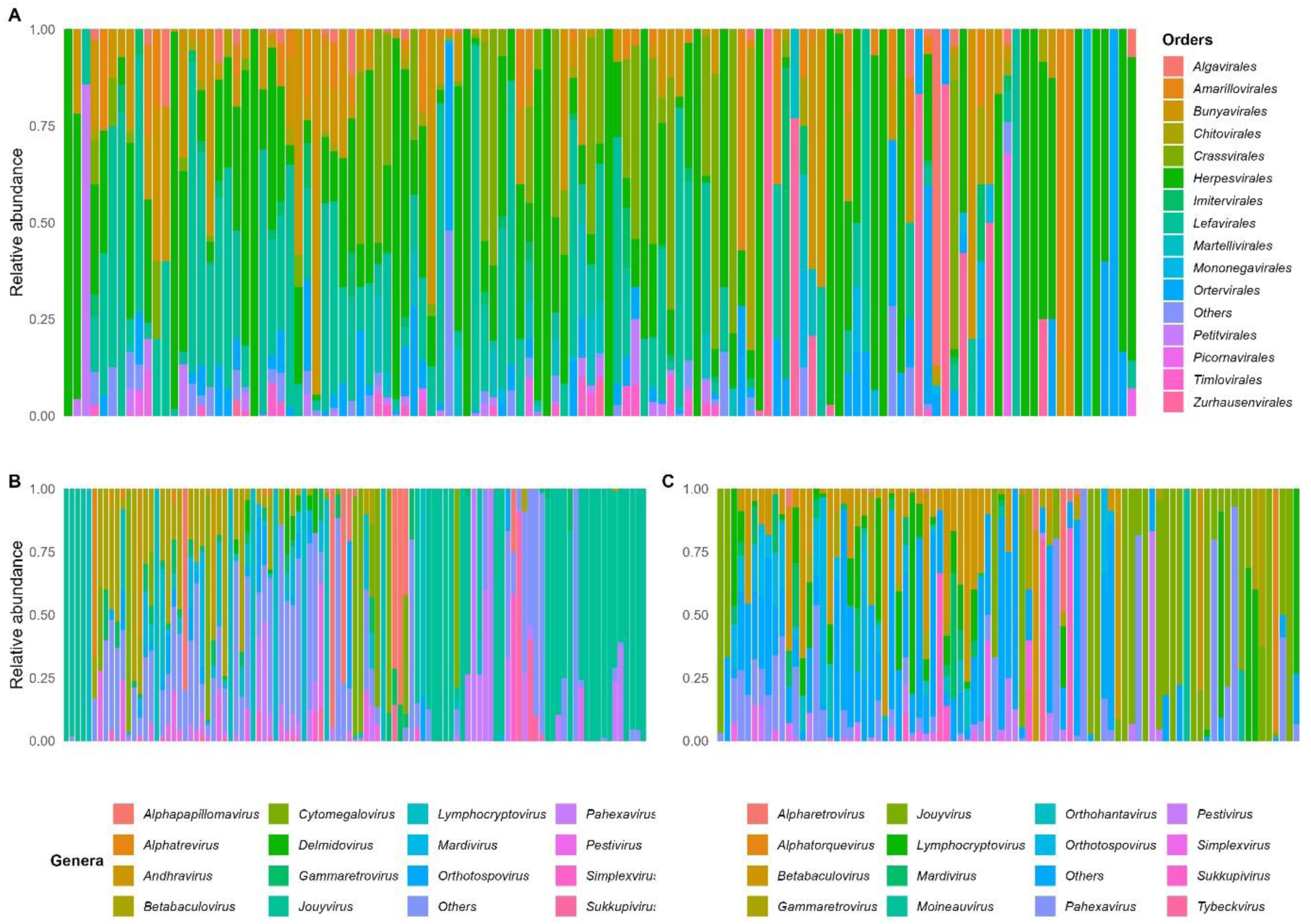
Taxonomic profile of the virome in gastric adenocarcinoma. (A) Most abundant viral orders overall. (B) Most abundant viral genera in tumor tissues. (C) Most abundant viral genera in peritumoral tissues.

### 3.2. Virome composition and abundance

Of the 329 viral genera, 101 were shared between the two tissues, while 109 were exclusive to tumor tissue and 119 were found only in peritumoral tissue (Figure 2A). To characterize the virome composition, the identified viral genera were classified based on their primary host into three categories: bacteriophages (viruses infecting bacteria), human viruses (viruses infecting humans), and other viruses (viruses infecting non-human eukaryotes, including animals, plants, protozoa, fungi and other invertebrates). In tumor tissue, 63.3% of the viral genera were bacteriophages, 7.6% were human viruses, and 29.1% were other viruses. A similar distribution was observed in peritumoral tissue, with 62.3% bacteriophages, 8.2% human viruses, and 29.5% other viruses (Figure 2B).

**Figure 2.**
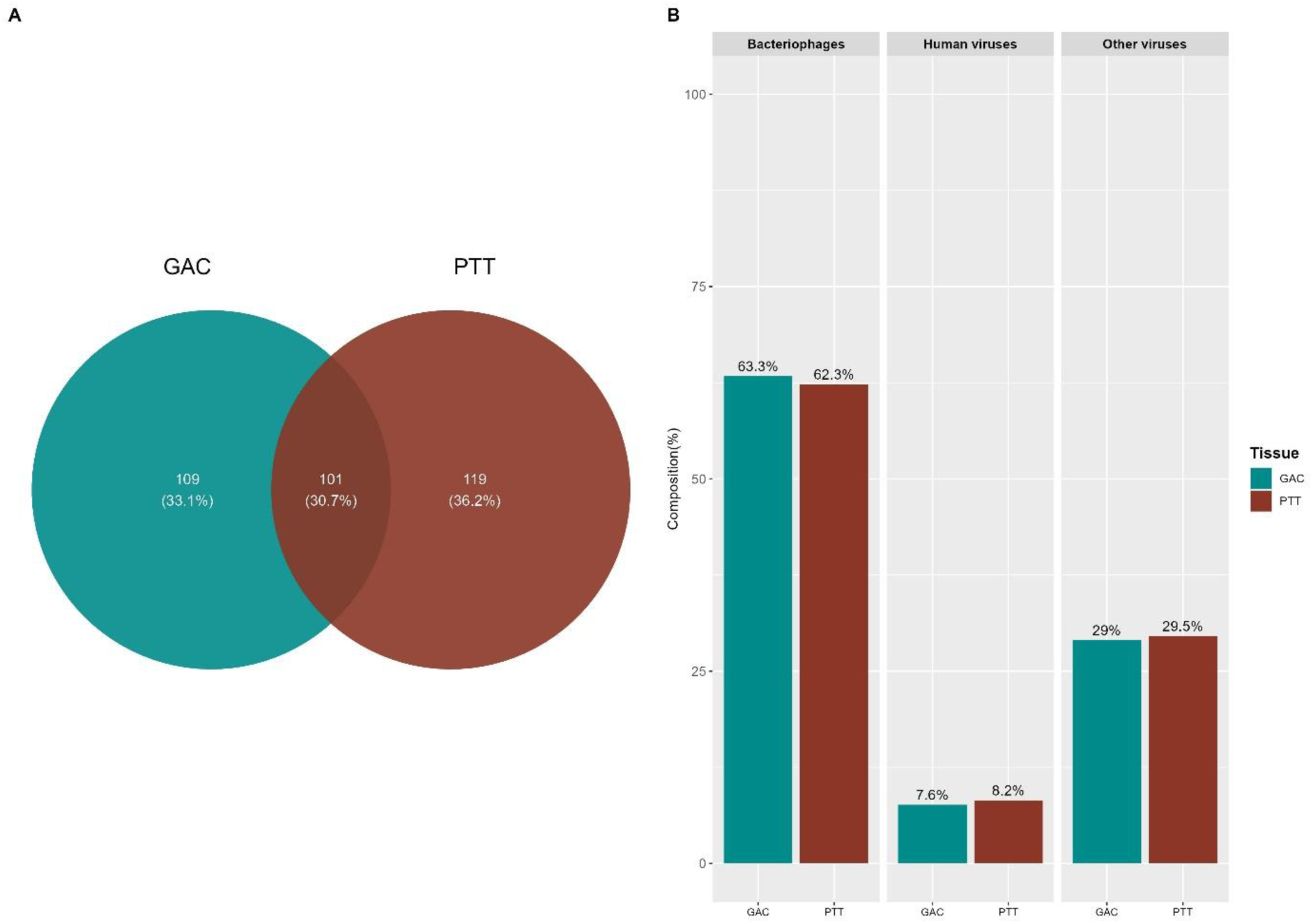
Virome composition in gastric adenocarcinoma. (A) Venn diagram showing unique and common viral genera between tumor and peritumoral tissues. (B) Classification of viral genera by host specifity.

A differential abundance analysis of the 101 viral genera shared between gastric tumor and peritumoral tissues was conducted using the Wilcoxon rank-sum test to identify significantly differentiated taxa. *Cytomegalovirus* (Wilcoxon, p = 0.003) and *Alphapapillomavirus* (Wilcoxon, p = 0.0133) were the most enriched in gastric tumors, whereas *Mastadenovirus* (Wilcoxon, p = 0.0013) and *Bahnicevirus* (Wilcoxon, p = 0.029) showed the most significant differences in gastric peritumoral tissues (Figure 3A-B).

**Figure 3.**
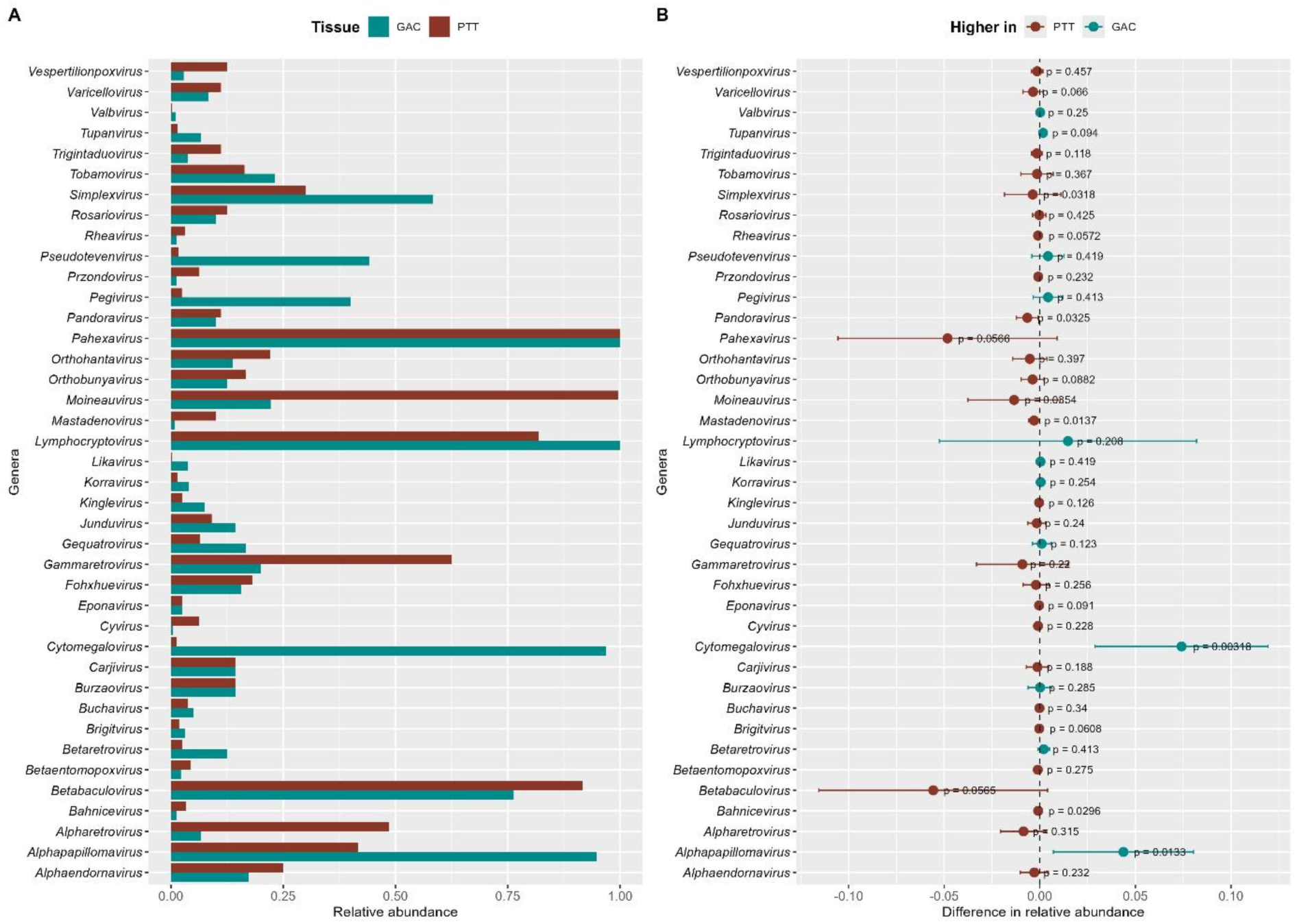
Differential viral abundance in gastric adenocarcinoma. (A) Viral genera differentially abundant in tumor and peritumoral tissues. (B) Mean differences in viral genera abundance between tumor and peritumoral tissues. Statistical comparisons were performed using the Wilcoxon test, with significance defined as p < 0.05. GAC - tumor tissue; PTT - peritumoral tissue.

### 3.3. Alpha and Beta diversity

Two metrics, alpha and beta diversity, were used to quantify the diversity of the viral community, assessing the impact of tissue type on viral diversity and composition in gastric tissues. Alpha diversity analysis showed no significant difference in observed richness between gastric tumor and peritumoral tissues (Wilcoxon, p = 0.16) (Figure 4A). For the Shannon index, a trend toward significance was observed (Wilcoxon, p = 0.0058), with peritumoral tissue exhibiting higher viral diversity (Figure 4B). The Simpson index also did not show a significant difference between tissues (WIlcoxon, p = 0.078) (Figure 4C).

**Figure 4.**
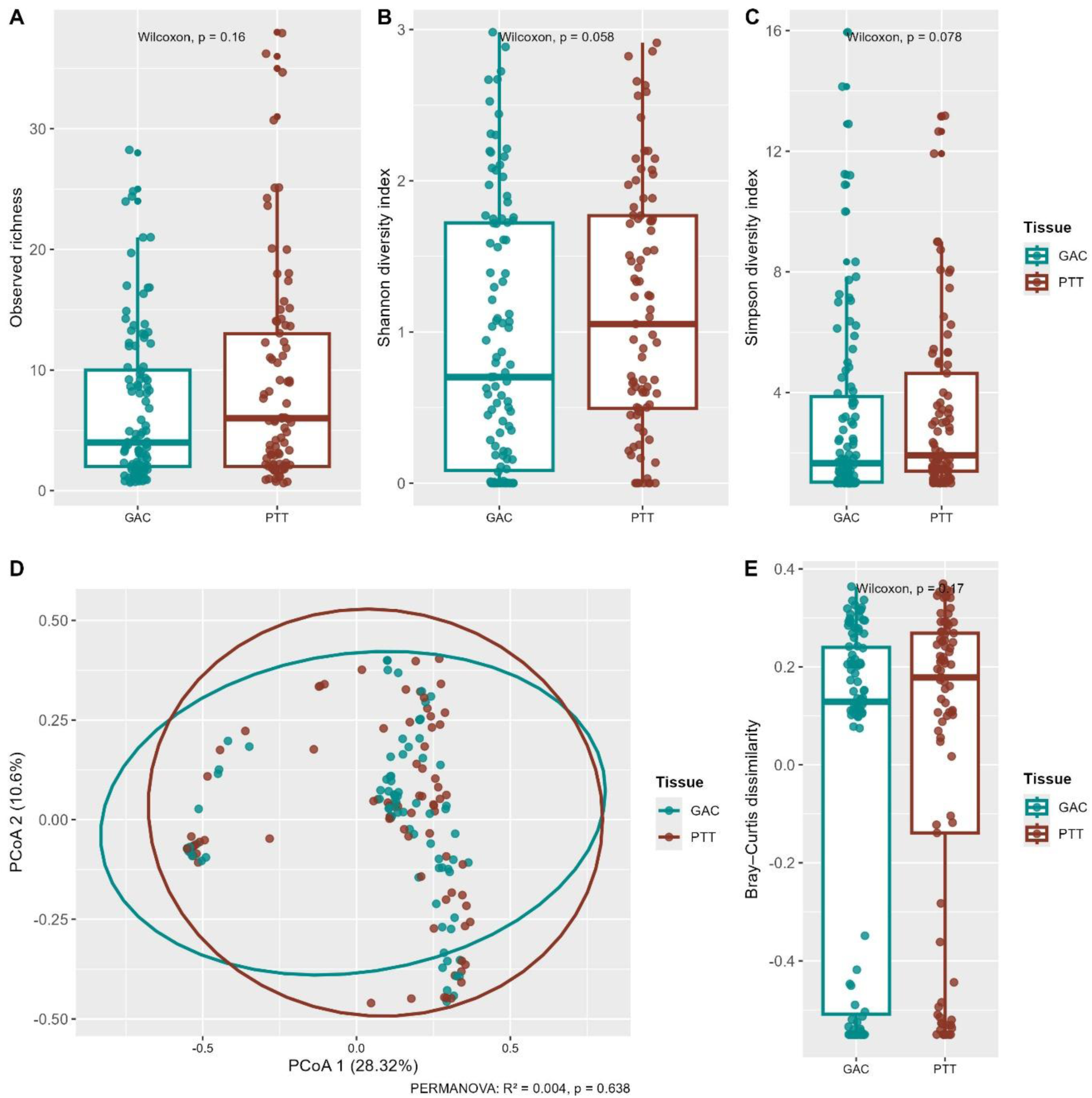
Alpha and Beta diversity in gastric adenocarcinoma. (A) Observed richness. (B) Shannon diversity index. (C) Simpson diversity index. (D) Principal Coordinate Analysis (PCoA) based on Bray–Curtis dissimilarity. (E) Bray–Curtis dissimilarity intra-group. Statistical comparisons were performed using PERMANOVA and the Wilcoxon test, with significance defined as p < 0.05. GAC - tumor tissue; PTT - peritumoral tissue.

Regarding the beta diversity analysis, the Bray-Curtis dissimilarity did not reveal a significant difference in viral composition (PERMANOVA, R² = 0.004, p = 0.629), indicating a similar viral taxonomic and abundance profile between gastric tumor and peritumoral tissues. PCoA1 explained 28.32% of the variation, and PCoA2 explained 10.6%, resulting in a cumulative explained variance of 38.92% across the samples (Figure 4D). Additionally, intra-group viral dissimilarity values also showed no significant differences between the tissues (Wilcoxon, p = 0.17) (Figure 4E).

## Discussion

Currently, several studies have investigated the role of microorganisms in the initiation and progression of gastric cancer. However, beyond *Helicobacter pylori*, the specific contribution of other components of the microbiota to gastric carcinogenesis has not been clearly established. Historically, gastric tissue was considered sterile due to its continuous exposure to a highly acidic environment. However, with the advancement of sequencing technologies, accumulating evidence has demonstrated that the stomach harbors a diverse microbial community^9, 20^.

In the present study, a total of 38 viral orders and 329 viral genera were detected in the analyzed gastric tissue samples. Among the most abundant viral orders, *Herpesvirales*, *Bunyavirales*, and *Ortervirales* stand out, known to include viruses capable of infecting a wide range of vertebrate hosts, including humans; *Lefavirales*, associated with invertebrate infections; and *Crassvirales*, encompassing the major families of bacteriophages^21–25^.

In gastric tumors, a total of 210 viral genera were identified, of which 109 were exclusive to this microenvironment. Regarding composition, 63.3% of the genera corresponded to bacteriophages, with *Jouyvirus*, *Pahexavirus*, *Sukkupivirus*, *Andhravirus*, *Delmidovirus*, and *Alphatrevirus* being the most abundant. Bacteriophages of the genera *Jouyvirus* and *Alphatrevirus* infect species of the genus *Escherichia*, commensals of the human gut, which have also been detected in gastric tissues^9, 26–28^.

*Pahexavirus* infect bacteria of the genera *Cutibacterium* and *Propionibacterium*, microorganisms commonly found on the skin that have also been detected in gastric tissues, which may explain the presence of these phages in the analyzed samples^29–32^. *Sukkupivirus* infect species of *Gordonia*, bacteria generally found in soil and water but capable of causing opportunistic infections in humans^33^. *Andhravirus* are *Staphylococcus* phages, a genus previously reported in gastric microbiomes, potentially representing transient colonization^34^. Finally, *Delmidovirus* is associated with *Bacteroides*, a genus previously identified in the gastric mucosa^35^.

It is widely recognized that phages do not directly infect human cells, exerting their influence indirectly on the host through the transfer of genetic material between bacteria, which can alter the fitness and virulence of these microorganisms. Additionally, prophage activation can result in bacterial lysis, thereby regulating the abundance of bacterial populations^36, 37^. Although the presence of phages in tumor gastric tissues does not directly imply a role in carcinogenesis, phage–bacteria interactions may modulate the local immune response, inflammation, and bacterial metabolite production, factors that could potentially contribute to tumor progression or the modulation of antitumor immunity^16, 38^.

In a study on colorectal cancer, a hypothetical mechanism termed “community-mediated viral oncogenesis” was proposed, in which certain bacteriophages may indirectly promote carcinogenesis. This process involves the lysis of commensal bacteria and the proliferation of opportunistic bacteria with carcinogenic potential. Concurrently, these opportunistic bacteria form expanding biofilms, in which phages play a key role by facilitating the invasion of the intestinal epithelium by oncogenic bacteria and contributing to tumorigenic transformation. These findings further support the indirect role of bacteriophages in tumor carcinogenesis^39^.

Viruses with tropism for human cells can establish latent infections, trigger immune responses, and, in certain cases, cause acute or chronic diseases^15^. In the present study, 7.6% of the viral genera identified in gastric tumors corresponded to viruses known to infect human cells, with the most abundant being *Lymphocryptovirus*, *Cytomegalovirus*, *Alphapapillomavirus*, and *Simplexvirus*, all widely recognized for their clinical relevance in human infections.

Although the abundance of *Lymphocryptovirus* did not differ significantly between tumor and peritumoral tissues in this study (Wilcoxon, p = 0.208), its presence in gastric tumors may have direct implications for carcinogenesis. *Lymphocryptovirus humangamma4*, traditionally known as *Epstein-Barr virus* (EBV), is the primary human representative of this genus and is associated with various hematological neoplasms, including Burkitt and Hodgkin lymphomas, as well as nasopharyngeal and gastric carcinomas^40–42^.

EBV-associated gastric tumors account for approximately 9% of gastric adenocarcinomas and represent a distinct subtype in terms of oncogenesis and molecular features, highlighting the importance of detecting this virus in gastric tumors^43^. EBV-related gastric carcinogenesis is primarily mediated by the expression of viral proteins, microRNAs (*BART*), and the BARF1 protein, which promote malignant transformation through the induction of methylation and the regulation of host gene expression^44^.

The representative species of *Cytomegalovirus* in the context of human infections is *Cytomegalovirus humanbeta5*, also known as Human cytomegalovirus (HCMV), one of the eight human herpesviruses, characterized by its ability to establish lifelong latency in the host and its potential for reactivation^45^. Although not officially classified as an oncogenic virus, HCMV infection is believed to contribute to the progression and development of various cancers, acting as an initiator or promoter of carcinogenesis, including breast, brain, and colorectal cancers^46–48^.

One study suggested that *Cytomegalovirus* infection may play a role in the progression or development of gastric cancer, possibly by modulating the tumor microenvironment in immunocompromised patients^49^. Furthermore, another study identified specific gene signatures associated with *Cytomegalovirus* infection in gastric cancer, indicating that the presence of the virus may influence tumor gene expression and, consequently, impact tumor progression^50^. In the present study, *Cytomegalovirus* exhibited significantly higher abundance in tumor tissues (Wilcoxon, p = 0.003); however, its exact role in gastric carcinogenesis remains to be fully elucidated.

*Alphapapillomavirus* infections, commonly referred to as Human papillomaviruses (HPVs), are highly prevalent among sexually active adults. Transmission occurs via direct contact with infected mucosal or cutaneous surfaces, and the viral genome may integrate into the host genome or persist in an episomal state^51^. HPV types 16 and 18 are the most common and are strongly linked to viral oncogenesis. The malignancies associated with these types primarily include cervical, penile, vulvar, vaginal, anal, and oropharyngeal carcinomas^52^.

Previous studies have reported the presence of high-risk HPVs in gastric adenocarcinomas, with detection in 5% (5/100) of samples in a study conducted in Iran^53^ and in 10 of 361 gastric tumor tissues in another study, of which 5 were genotyped as high-risk HPV-16, with active expression of the E6 and E7 oncogenes observed in only one sample^54^. In the present study, *Alphapapillomavirus* were significantly enriched in the analyzed gastric tumor tissues (Wilcoxon, p = 0.013). Although these viruses are traditionally associated with epithelial infections of the skin and oral and genital mucosae, their presence in gastric tissues is not uncommon, and their contribution to gastric carcinogenesis remains uncertain.

*Simplexvirus* constitutes a genus within the order *Herpesvirales*, with humans and other mammals serving as natural hosts^55^. The most prevalent species in humans are *Simplexvirus humanalpha1* and *Simplexvirus humanalpha2*, which are responsible for clinical manifestations such as oral herpes, genital herpes, herpetic stromal keratitis, erythema multiforme, herpetic eczema, meningitis, and encephalitis^56^. *Simplexvirus* sequences in gastric tissues is generally interpreted as a sequencing artifact, transient presence of viral particles from oral fluids, or contamination, rather than productive infection of the stomach.

Among the viral genera identified in gastric tumors in this study, 29.1% belonged to the group of “other viruses,” encompassing those that infect non-human eukaryotes, including animals, plants, protozoa, fungi, and other invertebrates. Among them, *Betabaculovirus* were relatively abundant. These viruses exclusively infect invertebrates and are widely used as biotechnological agents in the biological control of agricultural pests^57, 58^. Detection of *Betabaculovirus* in human gastric tissues may reflect accidental ingestion of food treated with bioinsecticides or, alternatively, misclassification of short or fragmented viral sequences that are homologous to *Betabaculovirus* but derive from other viruses or genetic elements of the gastric microbiota.

*Orthotospovirus* are plant pathogens that infect dicotyledonous crops of agricultural importance, particularly tomato (*Solanum lycopersicum*)^59^. Their detection in human gastric tissues is highly unlikely to represent a true infection, more likely reflecting the ingestion of contaminated food containing intact viral particles or RNA fragments. In contrast, *Gammaretrovirus* are well-recognized agents of leukemia in mammals such as cats, rodents, and non-human primates^60–62^. Although some exogenous *Gammaretrovirus* do not naturally infect humans, their endogenous elements (ERVs) are widely integrated into the human genome, which may explain their detection in the tumor samples analyzed in this study^63^.

*Pestivirus* are typically isolated from mammals such as cattle, in which they cause diarrhea and immunosuppression, and from swine, causing classical swine fever^64, 65^. *Mardivirus*, on the other hand, belong to the herpesviruses that primarily infect domestic birds and are responsible for economically significant diseases in poultry production^66^. The presence of these viruses in human gastric tissues does not indicate active infection, as their natural hosts are domestic mammals and birds, respectively. These findings likely reflect the consumption of contaminated animal meat and should be interpreted as environmental exposure or transient viral material. Alpha diversity quantifies the variety of taxa within a sample, considering both richness and evenness in the distribution of individuals^67^. In the present study, alpha diversity analysis revealed no significant differences between tumor and peritumoral tissues, both in observed richness (Wilcoxon, p = 0.16) and in Shannon (Wilcoxon, p = 0.0058) and Simpson indices (Wilcoxon, p = 0.078). These results indicate that viral richness and evenness are comparable between the two tissue types, suggesting that the presence of the tumor does not significantly alter intra-sample viral diversity.

Beta diversity quantifies differences in taxonomic composition between samples or communities, allowing the assessment of similarity or dissimilarity among groups^68^. Beta diversity analysis based on Bray-Curtis dissimilarity revealed no significant differences between tumor and peritumoral tissues in this study (PERMANOVA, R² = 0.004, p = 0.629), and the dissimilarity values among samples within each tissue group also did not differ significantly (Wilcoxon, p = 0.17). These results indicate that the viral composition is highly similar between the two tissue types, suggesting that the presence of the tumor does not significantly alter the local viral community.

## Conclusion

In the present study, we performed a characterization of the virome in tumor and peritumoral gastric tissues. Our results revealed a diverse viral community, comprising bacteriophages, human viruses, and other eukaryotic viruses, with distinct patterns of abundance between tumor and peritumoral tissues. While the overall viral diversity did not differ significantly between tissue types, specific human viruses, including *Lymphocryptovirus*, *Cytomegalovirus*, and *Alphapapillomavirus*, were enriched in tumor tissues, suggesting potential implications for gastric carcinogenesis. Bacteriophages were the predominant component, highlighting their possible indirect role in modulating bacterial populations, immune responses, and tumor microenvironment dynamics. These findings underscore the complexity of the gastric virome and provide a foundation for future investigations into the interactions and mechanisms through which the viral community could influence the development of gastric cancer, highlighting its potential role in gastric health and disease.

## Acknowledgment

The authors would like to thank the Oncology Research Center and the Human and Medical Genetics Laboratory at the Federal University of Pará (UFPA) for their invaluable technical and laboratory support. We are also grateful to the High-Performance Computing Center (CCAD) at UFPA for providing access to the Apollo 2000 cluster, which was essential for our analyses.

## Author contributions

**Conceptualization:** Diego Pereira, Fabiano Cordeiro Moreira, Paulo Pimentel de Assumpção, Rommel Mario Rodriguez Burbano. **Data curation:** Diego Pereira, Fabiano Cordeiro Moreira, Sérgio Augusto Antunes Ramos, Daniel de Souza Avelar. **Formal analysis:** Diego Pereira, Fabiano Cordeiro Moreira. **Funding acquisition:** Fabiano Cordeiro Moreira, Samia Demachki, Samir Mansour Casseb, Paulo Pimentel de Assumpção, Rommel Mario Rodriguez Burbano, Williams Fernandes Barra. **Investigation:** Diego Pereira, Fabiano Cordeiro Moreira, Daniel de Souza Avelar, Sérgio Augusto Antunes Ramos, Valéria Cristiane Santos da Silva, Jéssica Manoelli Costa da Silva, Ronald Matheus da Silva Mourão, Rubem Ferreira da Silva, Kauê Sant’Ana Pereira Guimarães, Juliana Barreto Albuquerque Pinto, Marcos da Conceição, Samir Mansour Casseb. **Methodology:** Diego Pereira, Fabiano Cordeiro Moreira, Daniel de Souza Avelar, Sérgio Augusto Antunes Ramos, Valéria Cristiane Santos da Silva, Jéssica Manoelli Costa da Silva, Ronald Matheus da Silva Mourão, Rubem Ferreira da Silva, Kauê Sant’Ana Pereira Guimarães, Juliana Barreto Albuquerque Pinto, Marcos da Conceição, Samir Mansour Casseb. **Sample acquisition:** Paulo Pimentel de Assumpção, Rommel Mario Rodriguez Burbano, Williams Fernandes Barra. **Supervision:** Fabiano Cordeiro Moreira, Samir Mansour Casseb, Paulo Pimentel de Assumpção, Rommel Mario Rodriguez Burbano. **Project administration:** Fabiano Cordeiro Moreira, Samia Demachki, Samir Mansour Casseb, Paulo Pimentel de Assumpção, Rommel Mario Rodriguez Burbano. **Visualization:** Diego Pereira, Fabiano Cordeiro Moreira. **Writing - original draft:** Diego Pereira, Fabiano Cordeiro Moreira. **Writing - review & editing:** Diego Pereira, Fabiano Cordeiro Moreira, Daniel de Souza Avelar, Sérgio Augusto Antunes Ramos, Valéria Cristiane Santos da Silva, Jéssica Manoelli Costa da Silva, Ronald Matheus da Silva Mourão, Rubem Ferreira da Silva, Kauê Sant’Ana Pereira Guimarães, Juliana Barreto Albuquerque Pinto, Marcos da Conceição, Williams Fernandes Barra, Samia Demachki, Samir Mansour Casseb, Rommel Mario Rodriguez Burbano, Paulo Pimentel de Assumpção.

## Data availability statement

All data generated or analyzed in this study are included within the article. Additional information is available from the corresponding author upon reasonable request.

## References

1. Thrift AP, El-Serag HB. Burden of Gastric Cancer. Clin Gastroenterol Hepatol. 2020;18(3):534–542. doi:10.1016/j.cgh.2019.07.045

2. Bray F, Laversanne M, Sung H, et al. Global cancer statistics 2022: GLOBOCAN estimates of incidence and mortality worldwide for 36 cancers in 185 countries. CA Cancer J Clin. 2024;74(3):229–263. doi:10.3322/caac.21834

3. Song Z, Wu Y, Yang J, Yang D, Fang X. Progress in the treatment of advanced gastric cancer. Tumour Biol J Int Soc Oncodevelopmental Biol Med. 2017;39(7):1010428317714626. doi:10.1177/1010428317714626

4. Wang B, Zhang Y, Qing T, et al. Comprehensive analysis of metastatic gastric cancer tumour cells using single-cell RNA-seq. Sci Rep. 2021;11(1):1141. doi:10.1038/s41598-020-80881-2

5. Yang L, Kartsonaki C, Yao P, et al. The relative and attributable risks of cardia and non-cardia gastric cancer associated with Helicobacter pylori infection in China: a case-cohort study. Lancet Public Health. 2021;6(12):e888–e896. doi:10.1016/S2468-2667(21)00164-X

6. Velanovich V, Hollingsworth J, Suresh P, Ben-Menachem T. Relationship of gastroesophageal reflux disease with adenocarcinoma of the distal esophagus and cardia. Dig Surg. 2002;19(5):349–353. doi:10.1159/000065835

7. Sexton RE, Al Hallak MN, Diab M, Azmi AS. Gastric cancer: a comprehensive review of current and future treatment strategies. Cancer Metastasis Rev. 2020;39(4):1179–1203. doi:10.1007/s10555-020-09925-3

8. Pereira-Marques J, Ferreira RM, Pinto-Ribeiro I, Figueiredo C. Helicobacter pylori Infection, the Gastric Microbiome and Gastric Cancer. Adv Exp Med Biol. 2019;1149:195–210. doi:10.1007/5584_2019_366

9. Ai B, Mei Y, Liang D, Wang T, Cai H, Yu D. Uncovering the special microbiota associated with occurrence and progression of gastric cancer by using RNA-sequencing. Sci Rep. 2023;13(1):5722. doi:10.1038/s41598-023-32809-9

10. Sahan AZ, Hazra TK, Das S. The Pivotal Role of DNA Repair in Infection Mediated-Inflammation and Cancer. Front Microbiol. 2018;9:663. doi:10.3389/fmicb.2018.00663

11. Wang Y, Han W, Wang N, et al. The role of microbiota in the development and treatment of gastric cancer. Front Oncol. 2023;13:1224669. doi:10.3389/fonc.2023.1224669

12. Gao JJ, Zhang Y, Gerhard M, et al. Association Between Gut Microbiota and Helicobacter pylori-Related Gastric Lesions in a High-Risk Population of Gastric Cancer. Front Cell Infect Microbiol. 2018;8:202. doi:10.3389/fcimb.2018.00202

13. Ferreira RM, Pereira-Marques J, Pinto-Ribeiro I, et al. Gastric microbial community profiling reveals a dysbiotic cancer-associated microbiota. Gut. 2018;67(2):226–236. doi:10.1136/gutjnl-2017-314205

14. Bakhti SZ, Latifi-Navid S. Interplay and cooperation of Helicobacter pylori and gut microbiota in gastric carcinogenesis. BMC Microbiol. 2021;21(1):258. doi:10.1186/s12866-021-02315-x

15. Liang G, Bushman FD. The human virome: assembly, composition and host interactions. Nat Rev Microbiol. 2021;19(8):514–527. doi:10.1038/s41579-021-00536-5

16. Broecker F, Moelling K. The Roles of the Virome in Cancer. Microorganisms. 2021;9(12):2538. doi:10.3390/microorganisms9122538

17. Zárate S, Taboada B, Yocupicio-Monroy M, Arias CF. Human Virome. Arch Med Res. 2017;48(8):701–716. doi:10.1016/j.arcmed.2018.01.005

18. Wood DE, Salzberg SL. Kraken: ultrafast metagenomic sequence classification using exact alignments. Genome Biol. 2014;15(3):R46. doi:10.1186/gb-2014-15-3-r46

19. Goodacre N, Aljanahi A, Nandakumar S, Mikailov M, Khan AS. A Reference Viral Database (RVDB) To Enhance Bioinformatics Analysis of High-Throughput Sequencing for Novel Virus Detection. mSphere. 2018;3(2):e00069–18. doi:10.1128/mSphereDirect.00069-18

20. Barra WF, Sarquis DP, Khayat AS, et al. Gastric Cancer Microbiome. Pathobiology. 2021;88(2):156–169. doi:10.1159/000512833

21. Dotto-Maurel A, Arzul I, Morga B, Chevignon G. Herpesviruses: overview of systematics, genomic complexity and life cycle. Virol J. 2025;22:155. doi:10.1186/s12985-025-02779-7

22. Boshra H. An Overview of the Infectious Cycle of Bunyaviruses. Viruses. 2022;14(10):2139. doi:10.3390/v14102139

23. Krupovic M, Blomberg J, Coffin JM, et al. Ortervirales: New Virus Order Unifying Five Families of Reverse-Transcribing Viruses. J Virol. 2018;92(12):e00515–18. doi:10.1128/JVI.00515-18

24. Wennmann JT. Nanopore reads spanning the whole genome of arthropod-infecting large dsDNA viruses of the class Naldaviricetes enable assembly-free sequence analysis. J Invertebr Pathol. 2025;209:108277. doi:10.1016/j.jip.2025.108277

25. Ramos-Barbero MD, Gómez-Gómez C, Sala-Comorera L, et al. Characterization of crAss-like phage isolates highlights Crassvirales genetic heterogeneity and worldwide distribution. Nat Commun. 2023;14:4295. doi:10.1038/s41467-023-40098-z

26. Mathieu A, Dion M, Deng L, et al. Virulent coliphages in 1-year-old children fecal samples are fewer, but more infectious than temperate coliphages. Nat Commun. 2020;11(1):378. doi:10.1038/s41467-019-14042-z

27. Rokyta DR, Burch CL, Caudle SB, Wichman HA. Horizontal gene transfer and the evolution of microvirid coliphage genomes. J Bacteriol. 2006;188(3):1134–1142. doi:10.1128/JB.188.3.1134-1142.2006

28. Hu YL, Pang W, Huang Y, Zhang Y, Zhang CJ. The Gastric Microbiome Is Perturbed in Advanced Gastric Adenocarcinoma Identified Through Shotgun Metagenomics. Front Cell Infect Microbiol. 2018;8:433. doi:10.3389/fcimb.2018.00433

29. Li X, Ding W, Li Z, et al. vB_CacS-HV1 as a Novel Pahexavirus Bacteriophage with Lytic and Anti-Biofilm Potential against Cutibacterium acnes. Microorganisms. 2024;12(8):1566. doi:10.3390/microorganisms12081566

30. Farrar MD, Howson KM, Bojar RA, et al. Genome Sequence and Analysis of a Propionibacterium acnes Bacteriophage. J Bacteriol. 2007;189(11):4161–4167. doi:10.1128/JB.00106-07

31. Lunger C, Shen Z, Holcombe H, et al. Gastric coinfection with thiopeptide-positive Cutibacterium acnes decreases FOXM1 and pro-inflammatory biomarker expression in a murine model of Helicobacter pylori-induced gastric cancer. Microbiol Spectr. 2024;12(1):e0345023. doi:10.1128/spectrum.03450-23

32. Li Q, Wu W, Gong D, Shang R, Wang J, Yu H. Propionibacterium acnes overabundance in gastric cancer promote M2 polarization of macrophages via a TLR4/PI3K/Akt signaling. Gastric Cancer Off J Int Gastric Cancer Assoc Jpn Gastric Cancer Assoc. 2021;24(6):1242–1253. doi:10.1007/s10120-021-01202-8

33. Mormeneo Bayo S, Palacián Ruíz MP, Asin Samper U, Millán Lou MI, Pascual Catalán A, Villuendas Usón MC. Pacemaker-induced endocarditis by Gordonia bronchialis. Enfermedades Infecc Microbiol Clin Engl Ed. 2022;40(5):255–257. doi:10.1016/j.eimce.2020.11.024

34. Shen Z, Dzink-Fox J, Feng Y, et al. Gastric Non-Helicobacter pylori Urease-Positive Staphylococcus epidermidis and Streptococcus salivarius Isolated from Humans Have Contrasting Effects on H. pylori-Associated Gastric Pathology and Host Immune Responses in a Murine Model of Gastric Cancer. mSphere. 7(1):e00772–21. doi:10.1128/msphere.00772-21

35. Herrera-Quintana L, Vázquez-Lorente H, Lopez-Garzon M, Cortés-Martín A, Plaza-Diaz J. Cancer and the Microbiome of the Human Body. Nutrients. 2024;16(16):2790. doi:10.3390/nu16162790

36. Taylor VL, Fitzpatrick AD, Islam Z, Maxwell KL. The Diverse Impacts of Phage Morons on Bacterial Fitness and Virulence. Adv Virus Res. 2019;103:1–31. doi:10.1016/bs.aivir.2018.08.001

37. Boling L, Cuevas DA, Grasis JA, et al. Dietary prophage inducers and antimicrobials: toward landscaping the human gut microbiome. Gut Microbes. 2020;11(4):721–734. doi:10.1080/19490976.2019.1701353

38. Cianci R, Caldarelli M, Brani P, et al. Cytokines Meet Phages: A Revolutionary Pathway to Modulating Immunity and Microbial Balance. Biomedicines. 2025;13(5):1202. doi:10.3390/biomedicines13051202

39. Hannigan GD, Duhaime MB, Ruffin MT, Koumpouras CC, Schloss PD. Diagnostic Potential and Interactive Dynamics of the Colorectal Cancer Virome. mBio. 2018;9(6):e02248–18. doi:10.1128/mBio.02248-18

40. Mui UN, Haley CT, Tyring SK. Viral Oncology: Molecular Biology and Pathogenesis. J Clin Med. 2017;6(12):111. doi:10.3390/jcm6120111

41. Pereira MA, Batista DAM, Ramos MFKP, et al. Epstein-Barr Virus Positive Gastric Cancer: A Distinct Subtype Candidate for Immunotherapy. J Surg Res. 2021;261:130–138. doi:10.1016/j.jss.2020.12.029

42. Li W, Duan X, Chen X, et al. Immunotherapeutic approaches in EBV-associated nasopharyngeal carcinoma. Front Immunol. 2022;13:1079515. doi:10.3389/fimmu.2022.1079515

43. Sunakawa Y, Lenz HJ. Molecular classification of gastric adenocarcinoma: translating new insights from the cancer genome atlas research network. Curr Treat Options Oncol. 2015;16(4):17. doi:10.1007/s11864-015-0331-y

44. Yang J, Liu Z, Zeng B, Hu G, Gan R. Epstein-Barr virus-associated gastric cancer: A distinct subtype. Cancer Lett. 2020;495:191–199. doi:10.1016/j.canlet.2020.09.019

45. Herbein G. Tumors and Cytomegalovirus: An Intimate Interplay. Viruses. 2022;14(4):812. doi:10.3390/v14040812

46. Herbein G, Kumar A. The Oncogenic Potential of Human Cytomegalovirus and Breast Cancer. Front Oncol. 2014;4:230. doi:10.3389/fonc.2014.00230

47. Peredo-Harvey I, Rahbar A, Söderberg-Nauclér C. Presence of the Human Cytomegalovirus in Glioblastomas—A Systematic Review. Cancers. 2021;13(20):5051. doi:10.3390/cancers13205051

48. Bai B, Wang X, Chen E, Zhu H. Human cytomegalovirus infection and colorectal cancer risk: a meta-analysis. Oncotarget. 2016;7(47):76735–76742. doi:10.18632/oncotarget.12523

49. Díaz-Alberola I, Espuch-Oliver A, Fernández-Segovia F, López-Nevot MÁ. Possible Role of Cytomegalovirus in Gastric Cancer Development and Recurrent Macrolide-Resistant Campylobacter jejuni Infection in Common Variable Immunodeficiency: A Case Report and Literature Discussion. Microorganisms. 2024;12(6):1078. doi:10.3390/microorganisms12061078

50. Krishnamoorthy P, Raj AS, Das N, et al. HCMV detection in Asian gastric cancer RNA-seq data sets and clinical validation in Indian GC patients reveals the HCMV-GC specific gene signatures. mSystems. 9(10):e00673–24. doi:10.1128/msystems.00673-24

51. Bogolyubova AV. [Human Oncogenic Viruses: Old Facts and New Hypotheses]. Mol Biol (Mosk). 2019;53(5):871–880. doi:10.1134/S0026898419050033

52. Szymonowicz KA, Chen J. Biological and clinical aspects of HPV-related cancers. Cancer Biol Med. 2020;17(4):864–878. doi:10.20892/j.issn.2095-3941.2020.0370

53. Fakhraei F, Haghshenas MR, Hosseini V, Rafiei A, Naghshvar F, Alizadeh-Navaei R. Detection of human papillomavirus DNA in gastric carcinoma specimens in a high-risk region of Iran. Biomed Rep. 2016;5(3):371–375. doi:10.3892/br.2016.728

54. Xu Q, Dong H, Wang Z, et al. Integration and viral oncogene expression of human papillomavirus type 16 in oropharyngeal squamous cell carcinoma and gastric cancer. J Med Virol. 2023;95(5):e28761. doi:10.1002/jmv.28761

55. Scott E, Burkhart C. Herpes simplex virus: An immunopathologic review. JAAD Rev. 2024;1:100–106. doi:10.1016/j.jdrv.2024.07.007

56. Tice B, Something J, Zimmerman B. Herpes Simplex Virus Infections: An Overview of Testing for the Urgent Care Clinician. JUCM J Urgent Care Med. 2025;19(9):31–36.

57. Shu R, Meng Q, Miao L, et al. Genome Analysis of a Novel Clade b Betabaculovirus Isolated from the Legume Pest Matsumuraeses phaseoli (Lepidoptera: Tortricidae). Viruses. 2020;12(10):1068. doi:10.3390/v12101068

58. Ferrelli ML, Salvador R. Effects of Mixed Baculovirus Infections in Biological Control: A Comprehensive Historical and Technical Analysis. Viruses. 2023;15(9):1838. doi:10.3390/v15091838

59. Kormelink R, Verchot J, Tao X, Desbiez C. The Bunyavirales: The Plant-Infecting Counterparts. Viruses. 2021;13(5):842. doi:10.3390/v13050842

60. Hofmann-Lehmann R, Hartmann K. Feline leukaemia virus infection: A practical approach to diagnosis. J Feline Med Surg. 2020;22(9):831–846. doi:10.1177/1098612X20941785

61. Akkawi C, Feuillard J, Diaz FL, et al. Murine leukemia virus (MLV) P50 protein induces cell transformation via transcriptional regulatory function. Retrovirology. 2023;20(1):16. doi:10.1186/s12977-023-00631-w

62. McKee J, Clark N, Shapter F, Simmons G. A new look at the origins of gibbon ape leukemia virus. Virus Genes. 2017;53(2):165–172. doi:10.1007/s11262-017-1436-0

63. Bao C, Gao Q, Xiang H, et al. Human endogenous retroviruses and exogenous viral infections. Front Cell Infect Microbiol. 2024;14:1439292. doi:10.3389/fcimb.2024.1439292

64. Wernike K, Gethmann J, Pfaff F, Sauter-Louis C, Beer M. Bovine viral diarrhea virus eradication in Germany: A never-ending success story or just the last 46 PI animals? Vet Microbiol. 2025;309:110697. doi:10.1016/j.vetmic.2025.110697

65. Hochman O, Goonewardene K, Chung CJ, Ambagala A. Evaluation of Spleen Swabs for Sensitive and High-Throughput Detection of Classical Swine Fever Virus. Pathog Basel Switz. 2025;14(8):767. doi:10.3390/pathogens14080767

66. Durand M, Chuard A, Rémy S, et al. The pUL47 tegument protein of Marek’s Disease Virus interacts with p32/C1QBP to promote horizontal transmission. PLoS Pathog. 2025;21(8):e1013434. doi:10.1371/journal.ppat.1013434

67. Thukral AK. A review on measurement of Alpha diversity in biology. Agric Res J. 2017;54(1):1. doi:10.5958/2395-146X.2017.00001.1

68. Tuomisto H. A diversity of beta diversities: straightening up a concept gone awry. Part 1. Defining beta diversity as a function of alpha and gamma diversity. Ecography. 2010;33(1):2–22. doi:10.1111/j.1600-0587.2009.05880.x

